# Black pod disease profile: Monitoring its outbreak in Southwest, Nigeria

**DOI:** 10.1101/479501

**Authors:** Peter M. Etaware, Adegboyega R. Adedeji

## Abstract

Black pod disease (BPD) has been and still remains a major threat to cocoa farmers worldwide due to its annual recurrence, fast spread and highly destructive nature. The disease has caused great anxiety in many cocoa producing communities due to the inability of indigenous cocoa farmers to determine when and where BPD outbreak will take place. Twelve (12) stations were structured from four important cocoa-producing States in the Southwestern region of Nigeria. An investigation of BPD outbreak was conducted in 2015/2016 within these regions. Infected cocoa pods and topsoil samples were collected for laboratory analysis. Pests attack, cherelle wilt and BPD outbreak were seasonal with 50% chances of occurrence in all the stations. Black pod diseases outbreak was recorded in all the States (100%) during the rainy season. The disease was at its peak in August 2015 in almost all the stations (station 1 (30.0%), station 3 (23.0%), station 11 (16.0%), station 4 (9.0%), station 5 (7.0%), and station 8 (3.0%). The height of disease severity was in September 2015 (station 1 (100.0%), station 3 (96.7%), station 5 (85.7%), station 11 (84.3%), and station 4 (70.0%), with station 8 reaching the 100% mark in October 2015. Most cocoa farmlands are now being abandoned, unless concerted efforts are made to effectively manage the disease, BPD will greatly reduce cocoa production in Nigeria and around the world.

## Introduction

*Theobroma cacao* Linn. (Cocoa) is an evergreen and relatively common understory tree growing 4 - 8 m tall. Cocoa is native to the deep tropical region of South America, and is now widely cultivated in Africa and some parts of Europe, Asia and Australia. Cocoa cultivation has been a major source of income for most third world countries. The largest proportion of global cocoa beans (59%) comes from West Africa, with Côte d’Ivoire and Ghana producing 1,472,313 858,720 tonnes respectively in 2016 [1]. Unlike many exotic crops, Cocoa is highly susceptible to pestilence [2]. These pestilences occur seasonally and as such are influenced by weather, they mostly affect coupons, cherelles, unripe and ripe cocoa pods at different developmental stages in the field [3].

The earliest account of cocoa pestilence in Africa was reported by Thorold [4]. The topmost invasive diseases of cocoa are black pod disease, swollen shoot, witches’ broom, *Monilia* pod rot, and vascular-streak dieback [3]. Black pod disease has been the most recurrent, highly destructive, dreaded and widespread among other diseases [3]. This disease however seems more established in West Africa than in other cocoa-growing regions around the world. In Nigeria, Cameroon and Togo, black pod disease constitutes the greatest set-back to cocoa production with losses of up to 90% [3]. In the early 1980s, black pod disease in Ghana was only known to be caused by *P. Palmivora.* However, in 1985, a severe outbreak, which appeared different from that previously known, was reported in the Akomadan area of the Ashanti region. Laboratory investigations by Dakwa [5] on diseased cocoa pod samples showed that *P. megakarya* was the causal agent, and this was subsequently confirmed by several researchers [6,7]. This was the first reported incidence of the species in Ghana, but earlier observations and research activities carried out in the Volta region indicated that the species might have existed there, perhaps as far back as the early 1970s. *Phytophthora megakarya* had been reported in many other African nations including Gabon, Equatorial Guinea where it was reputed for massive declines in cocoa production before 1985, [3].

The occurrence of *P. megakarya* in West Africa has changed the status of black pod disease in Ghana, Nigeria, Cameroun and other regions involved in cocoa production. Black pod disease has been attributed to several species *of Phytophthora* such as *P. drechsleri* [8], *P. botryosa* [9],

*P. heveae* [10], *P. meadii* [11], *P. capsici* [12], *P. megakarya* [13], *P. citrophthora* [14], *P. katsurae* [15] and *P. tropicalis* [16]. Black pod disease caused by *P. megakarya* continues to be the major threat to cocoa production in West Africa [3,6]. Virulence in *P. megakarya* emanates from the ability to produce large number of spores on the pod surface [17].

*P. megakarya* produces lesions of black pod disease with irregular edges on the pod surface whereas lesions caused by *P. palmivora* have regular borders and are generally smaller [18]. The first symptom is a brown to black spot on the pod, which spreads rapidly in all directions and eventually covers the whole pod. The beans become infected internally about 15 days after the initial infection and are soon of no commercial value [19]. Generally, pods closest to the ground are first infected, with the disease rapidly spreading to affect fruit on the entire tree. *Phytophthora megakarya* can also cause seedling blight and trunk cankers [20]

The disease now poses a serious threat to the cocoa industry and has caused great anxiety and concern in many cocoa producing communities. Pod losses to *P. megakarya* are massive and some farmers in the affected areas have had virtually no crop for many seasons. As a result of the disease, cocoa farms are being neglected or totally abandonedand some farmers seemingly have little or no enthusiasm in establishing new cocoa farms in the areas where the percentage of black pod disease occurrence is very high [3]. In addition, most farm concierges have turned their attention to other crops due to the incessant black pod disease infestation on cocoa crop. Reports from indigenous cocoa farmers and extension workers as well as reports from the field during black pod surveys indicate that *P. megakarya* proliferates faster and it has spread to some important cocoa-growing areas leading to an upsurge of the disease. The disease is not only responsible for immense pod losses, but also infliction of severe stem canker resulting in the death of many cocoa trees [7].

The success rate achieved by both biological and chemical control measures is fast declining due to the high level of adaptation of the pathogen to harsh conditions, the constant change in rainfall pattern and irregular fluctuation of other weather parameters coupled with the drastic increase in vulnerability of cocoa plants. The dearth of information on the ontogeny and phylogenetic trend of the species of *Phytophthora* involved in the recurrent annual perpetuation of black pod disease in cocoa producing areas within the rural and sub-urban communities in Nigeria is a major setback to the effective management of the disease in Nigeria. Hence, there is an urgent need for a revolutionized approach in the management of black pod disease. Unless concerted efforts are made to effectively manage the disease BPD will greatly reduce cocoa production in Nigeria and around the world [3,21]. Therefore, this study was designed to develop a system for black pod disease prediction in order to provide useful and timely information on black pod disease occurrence, its severity and the specific areas expected to be affected. This will minimize fungicide misuse, increase cocoa productivity, reduce the risk of chemical poisoning, increase farmers’ profit, foreign exchange and internally generated revenue, and ensure the availability of disease-free and non-toxic raw materials for cocoa processing industries. Lastly, it will create a clean and healthy environment for the sustenance of all forms of life.

## Materials and methods

### Research Locations

Twelve (12) commercial cocoa farms in Southwest, Nigeria were selected for black pod disease assessment (Table 1; S1 Appendix). The criteria for selection were: 1) farm size (The minimum acceptable size was 10,000 m^2^), 2) consistency in cocoa production, 3) cropping system and 4) locality. These were used to determine the suitability of the selected cocoa farms in line with the aim of the current research. The selected locations were classed as stations for ease of identification (Figs 1 - 2).

**Table 1:** Location of sampled stations and size of farms surveyed.

| Station | Location | Latitude N | Longitude E | Altitude (m) | Size (m<sup>2</sup>) |
| --- | --- | --- | --- | --- | --- |
| 1 | Òwenà | 7° 12' 11.52" | 5° 00' 55.76" | 289 | 10,000 |
| 2 | Òwenà | 7° 12' 11.50" | 5° 00' 55.76" | 291 | 10,000 |
| 3 | Wáàsimi | 7° 10' 42.78" | 4° 59' 31.34" | 249 | 30,000 |
| 4 | Adaàgbà | 7° 22' 13.80" | 4° 33' 34.42" | 262 | 40,000 |
| 5 | Iyánfowórogì | 7° 21' 55.22" | 4° 34' 16.54" | 259 | 20,000 |
| 6 | Owódé-Igàngán | 7° 29' 59.99" | 4° 48' 59.99" | 276 | 50,000 |
| 7 | Owódé-Igàngán | 7° 29' 53.45" | 4° 48' 59.01" | 282 | 50,000 |
| 8 | Qbáfémi-Owódé | 7° 08' 30.37" | 3° 25' 56.71" | 187 | 10,000 |
| 9 | Qbáfémi-Owódé | 7° 08' 30.32' | 3° 25' 56.73" | 192 | 10,000 |
| 10 | Mòyè | 7° 18' 54.54" | 4° 01' 09.34" | 205 | 20,000 |
| 11 | Qmi-Adió | 7° 20' 47.58" | 3° 44' 30.59" | 174 | 20,000 |
| 12 | Olórò | 7° 20' 44.00" | 3° 59' 34.00" | 179 | 10,000 |

**Fig 1:**
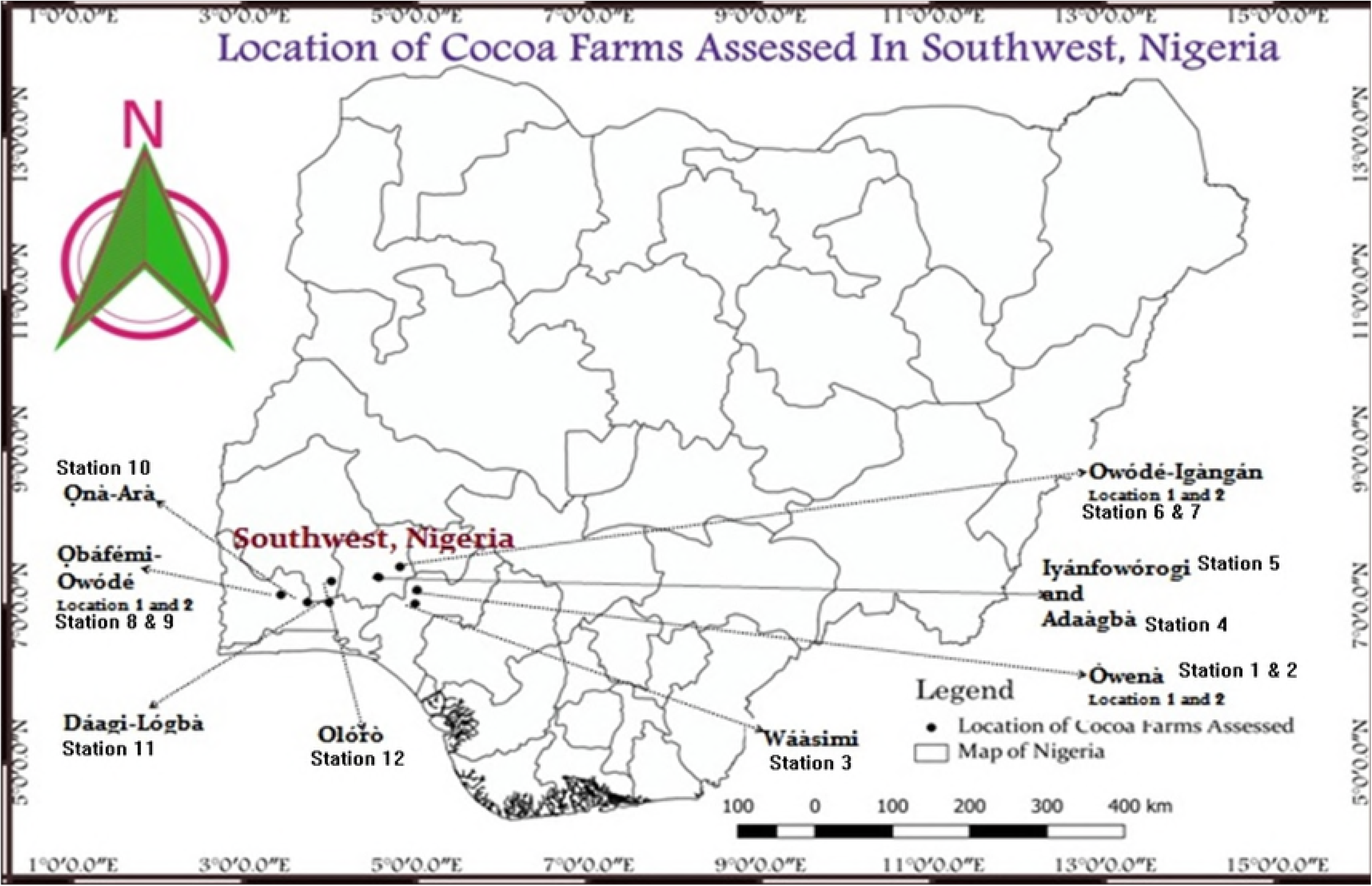
The geographical locations of the selected cocoa farm stations in Soutwest Nigeria.

### Epidemiological Survey within the Stations

Black pod disease assessment was conducted for thirteen (13) consecutive months. The period of the disease assessment covered the major cocoa growing season (March to October), the minor cocoa production season (November to April) and the optimum cocoa growing season (July to August) for cocoa production in Nigeria. The minimum Cocoa farm size considered for the assessment of black pod disease incidence and severity was ten thousand square meters (10,000 m^2^) or one hectare (1 hectare).

The disease assessment was conducted both in the rainy season and dry season to determine the level of variation of black pod disease outbreak and severity brought about by seasonal changes. Also, the altitude of the study areas was considered in other to determine its influence on disease development and spread. Therefore, the study locations were classed accordingly and the observations made were grouped based on the established criterion.

### Black Pod Disease Occurrence

The method adapted for black pod disease incidence determination was that of Luo [22]. Cocoa trees were assessed in a transverse and diagonal mode as described in Fig 2 and Plate 4 within each study location. Green and ripe Cocoa pods from each tree were inspected for the symptoms of black pod disease; the rain splash zone described in Plate 5 was of interest. If an infected pod was detected on the tree, the stand (tree) was noted as being infected. The assessment was repeated for a total of one hundred (100) trees and the observations noted. Each tree stand was noted as disease free (Healthy) or Infected based on the presence or absence of black pod disease symptoms. The observations were carried out for thirteen (13) months (May, 2015 to May, 2016).

**Fig 2:**
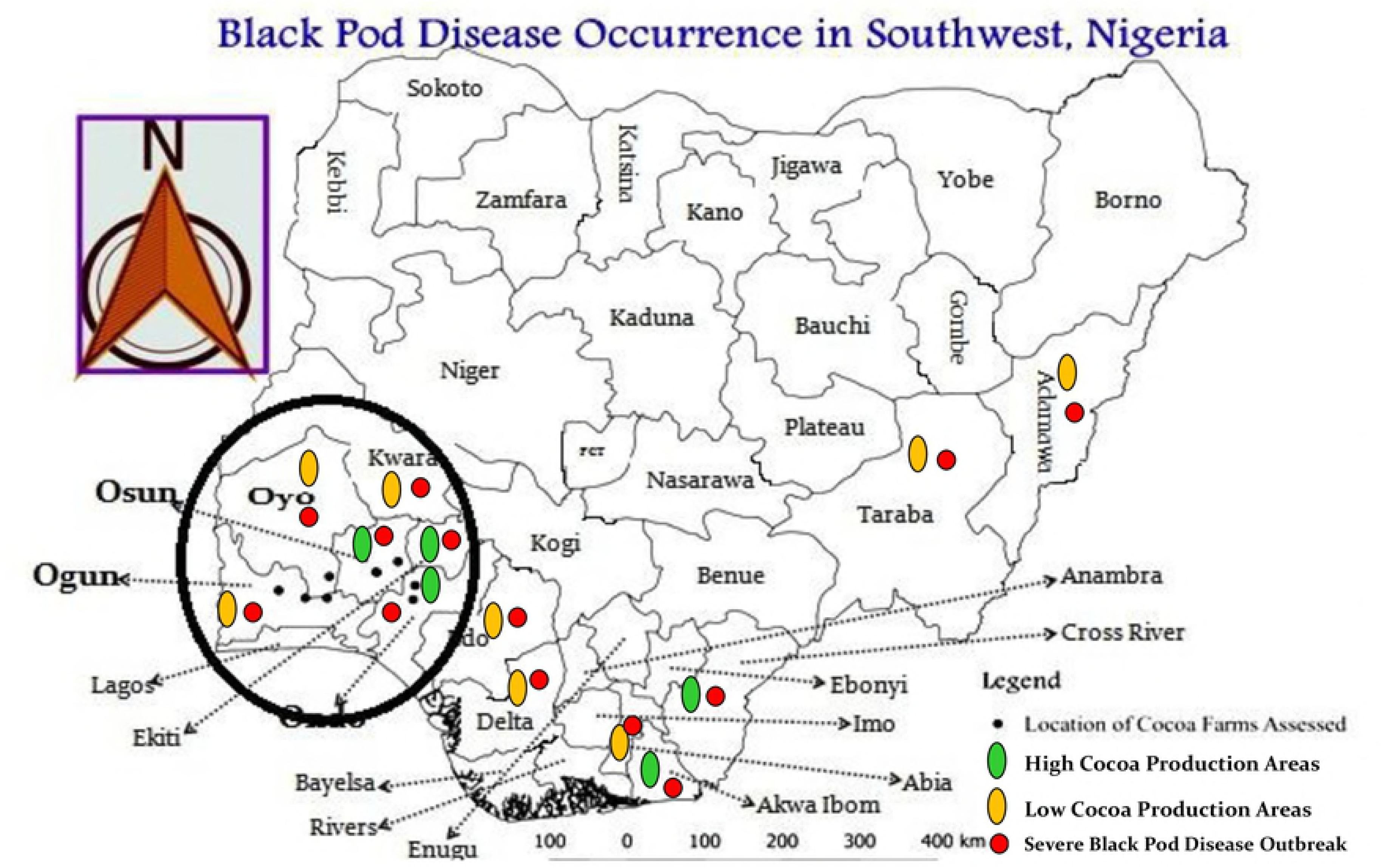
The severity intensity of black pod disease in Nigeria (2015/2016)

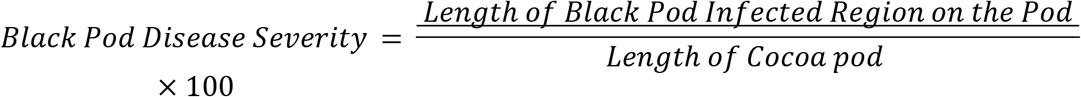

### Black Pod Disease Severity

For the determination of black pod disease severity, Cocoa trees were also assessed in a transverse and diagonal mode as shown in S1 Fig [23]. Two methods were adapted for black pod disease severity determination to minimize error(s). The first method involves the measurement of infected cocoa pods and the percentage infection calculated based on the total Cocoa pod length basically from the rain splash zone (Fig 3). The second method involves the superficial assessment of the extent of damage inflicted by the disease on each infected cocoa pod and a score from 0 to 5 ascribed (Table 2) [24]. This scored served as the rating for the disease infection for that particular cocoa pod.

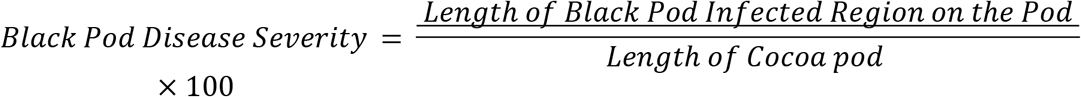

**Table 2:** Black pod disease severity status determination

| **BPD INFECTION per COCOA POD (BPD Severity)** |  |  |  |
| --- | --- | --- | --- |
| **Score** | **Infected Portion** | **Percentage Infection** | **Inference** |
| 0 | None | 0% | Healthy |
| 1 | $(1/5)$ | 20% | Not Severe |
| 2 | $(2/5)$ | 40% | Mildly Severe |
| 3 | $(3/5)$ | 60% | Averagely Severe |
| 4 | $(4/5)$ | 80% | Severe |
| 5 | All | 100% | Extremely Severe |

An Adaptation of the modified methods of Akrofi [24]. BPD – Black pod disease

**Fig 3:**
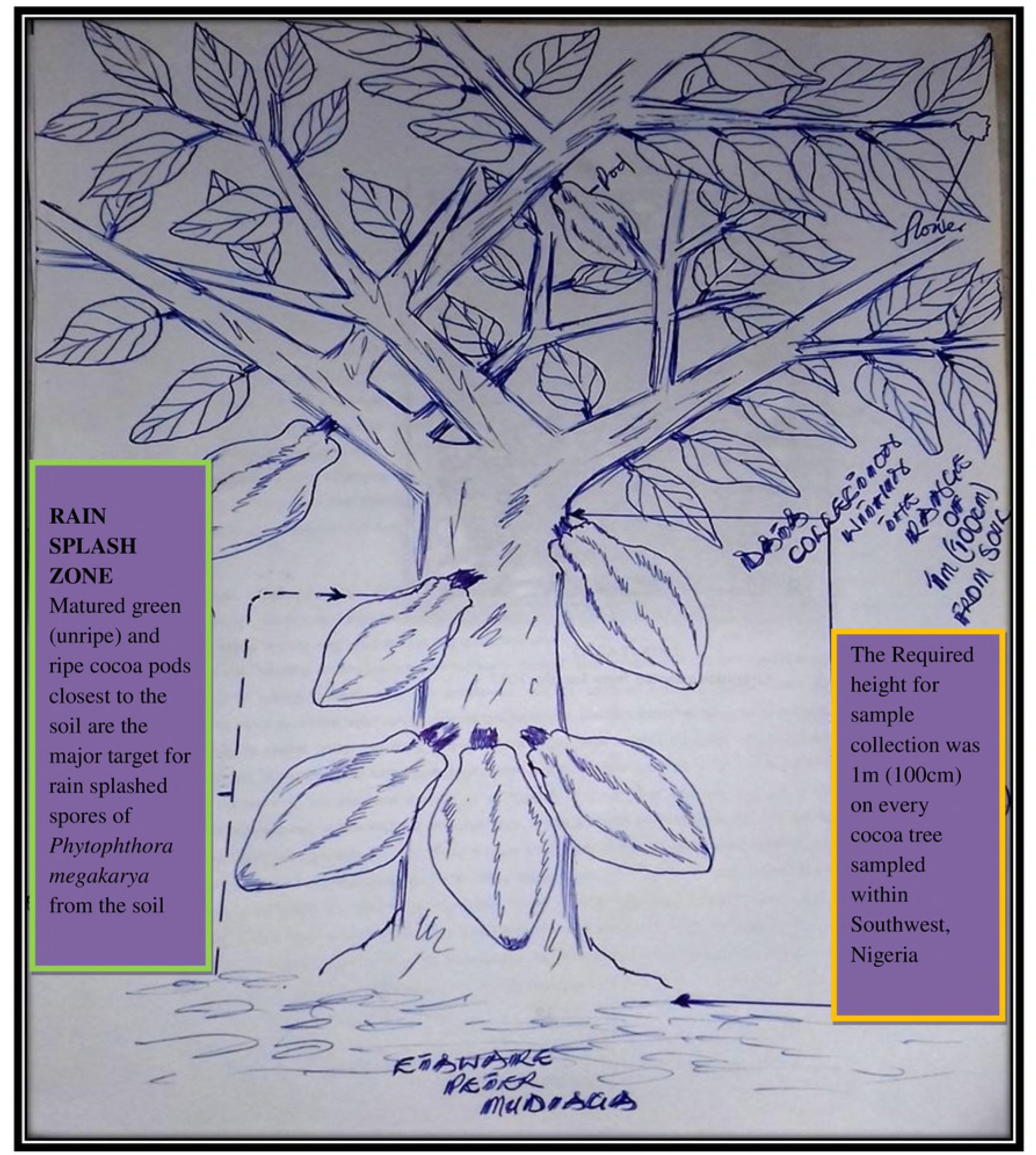
The mode of biological data collection on the field

In general, the observations made within each Season, Station Altitude, State and the Southwest in general were calculated using the following formulae:

Observations recorded per Season: % SI = (SOS_i_ /SL_total_)

Where SI = Seasonal Influence on BPD Status, SOS_i_ = Sum of all observations made per Season (per month) and SL_total_ = The Total No. of study locations assessed

Observations recorded per Altitude level: % AI = (SOA_i_ /SL_total_)

Where AI = Altitudinal influence on BPD Status, SOA_i_ = Sum of all observations made per altitudinal Level (per month) and SL_total_ = The Total No. of study locations assessed

BPD Status in Southwest Nigeria: % BSS = (SO_i_ /SL_total_)

Where BSS = BPD Status in Southwest, Nigeria, SO_i_ = Sum of all observations (per month) from the stations and SL_total_ = The Total No. of study locations assessed

For BPD Status in States: % BST = (ST_i_ /SL_total_)

Where BST = BPD Status in each State, SOS_i_ = the sum of BPD Status from all the sampled stations within a State (per month) and SL_total_ = The Total No. of study locations assessed

### Previous black pod disease records

The previous disease record on the prevalence and intensity of black pod disease of cocoa was obtained from Cocoa Research Institute of Nigeria (CRIN), Idi-Ayunre, Ibadan, Oyo State, Nigeria and the report of Lawal and Emaku [23]. The data collected spanned from 1985 to 2014.

### Data Analysis

Data were collected from the rain splash zone, root rhizosphere and husk dumpsite as shown in Fig 3. Qualitative data were represented as charts and graphs plotted using Microsoft Office (Excel) 2007 service pack and SPSS version 20.0 for 32 bits resolution. An Analysis of variance was carried out using COSTAT 9.0 software, while the homogeneity of means was determined using Duncan Multiple Range Test (DMRT).

## Results

### Orientation on PDA (Morphological Characters)

The pathogen appeared cotton white on potato dextrose agar (PDA) from day one. A single cluster of the velvet mycelia colony was formed around the Petri-plate. The silky mycelia colony turned pale creamy-yellow in older cultures due to the production of light lemon-yellowish secretion which could be a secondary metabolite or an extracellular secreted enzyme (no further test was conducted to determine the nature of the metabolite secreted) around the midpoint of inoculation as shown in Fig 4a

**Fig 4:**
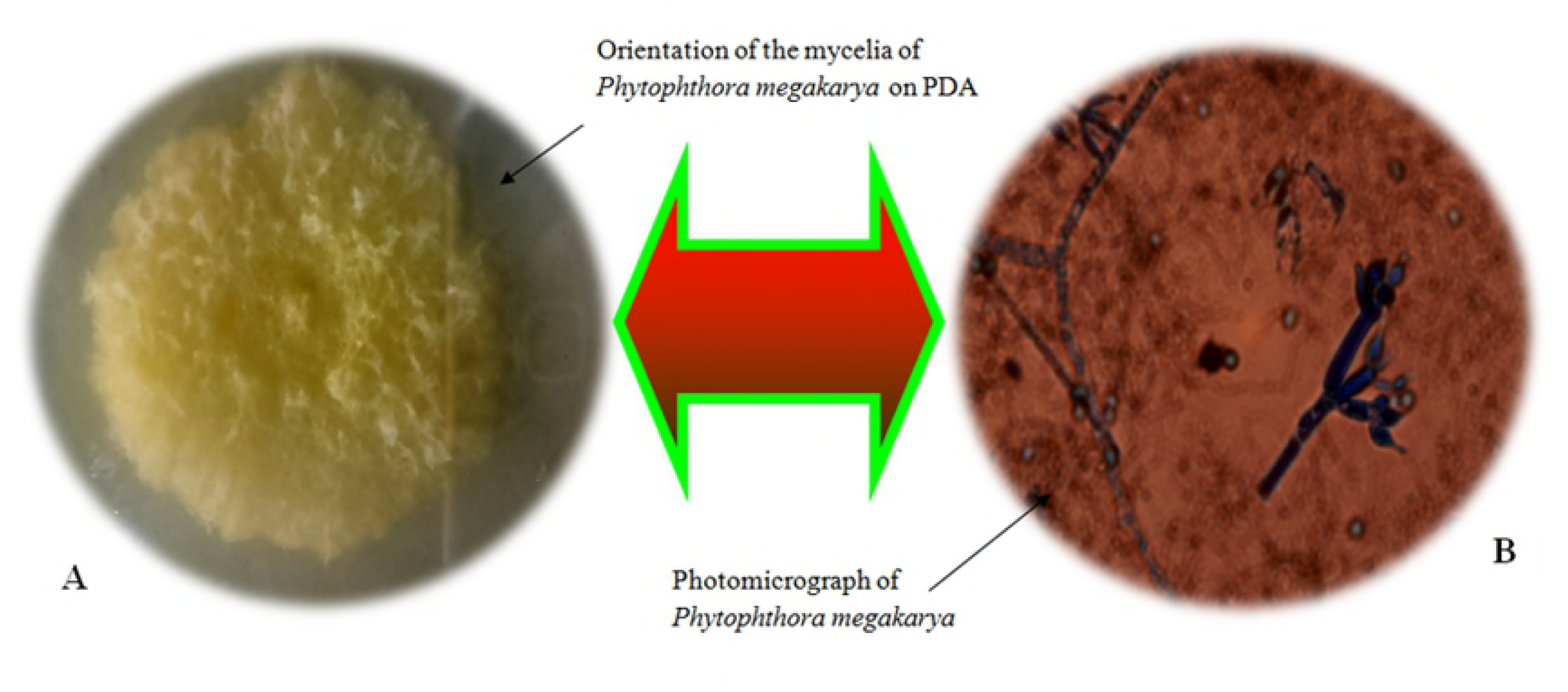
The morphological and cytological features of *Phytophthora megakarya* on potato dextrose agar (PDA) and under the microscope

### Microscopy (Cytological Characters)

The type of spore(s) produced i.e. chlamydospores (Zoospores), Oospores (Sexual spores) or aplanospores was the distinguishing factor within the various species of *Phytophthora* notable for inciting black pod disease. Micro-images of the hyphal structure appeared hyaline, septate and heterogeneously branched, double walled with thin layers. The production of zoospores on special reproductive hyphae known as the sporangiophore was also noticed. The zoospores produced were double layered, ellipsoidal/oval in shape, with a pointed node each for attachment to the sporangiophore. Each zoospore had a single flagellum that facilitated mobility. The flagellum was short and located at the posterior part of the spore. The spore stained purple to violet when exposed to lactophenol in cotton blue dye and were categorically attached singly at the apices of the sporangiophore (Fig 4b).

### The life cycle of *Phytophthora megakarya*

The life cycle of *P. megakarya* was studied in the Mycology/Pathology Laboratory of the Department of Botany, University of Ibadan (Fig 5). Spores of the pathogen germinated into motile zoospores (Fig 5b). The motile spores lose their flagella, germinate, and produce infection peg alongside penetration mechanisms in order to gain access into the host tissue (Fig 5c). After germination, the pathogen produces somatic cell structures, mycelia and hyphal structures (Fig 5d). At maturity, the pathogen produces fruiting structures (sporangiophores) with spores apically located at the end of the hyphae (Fig 5e). Sporangiophores of *Phytophthora megakarya* are branched reproductive hyphae with thick walls. The sporangiophore bears spores that are attached to it by the peduncle (Fig 5e). Mature flagellated zoospores are then released into the environment when there is distress or limitation in food supply (Fig 5a).

**Fig 5:**
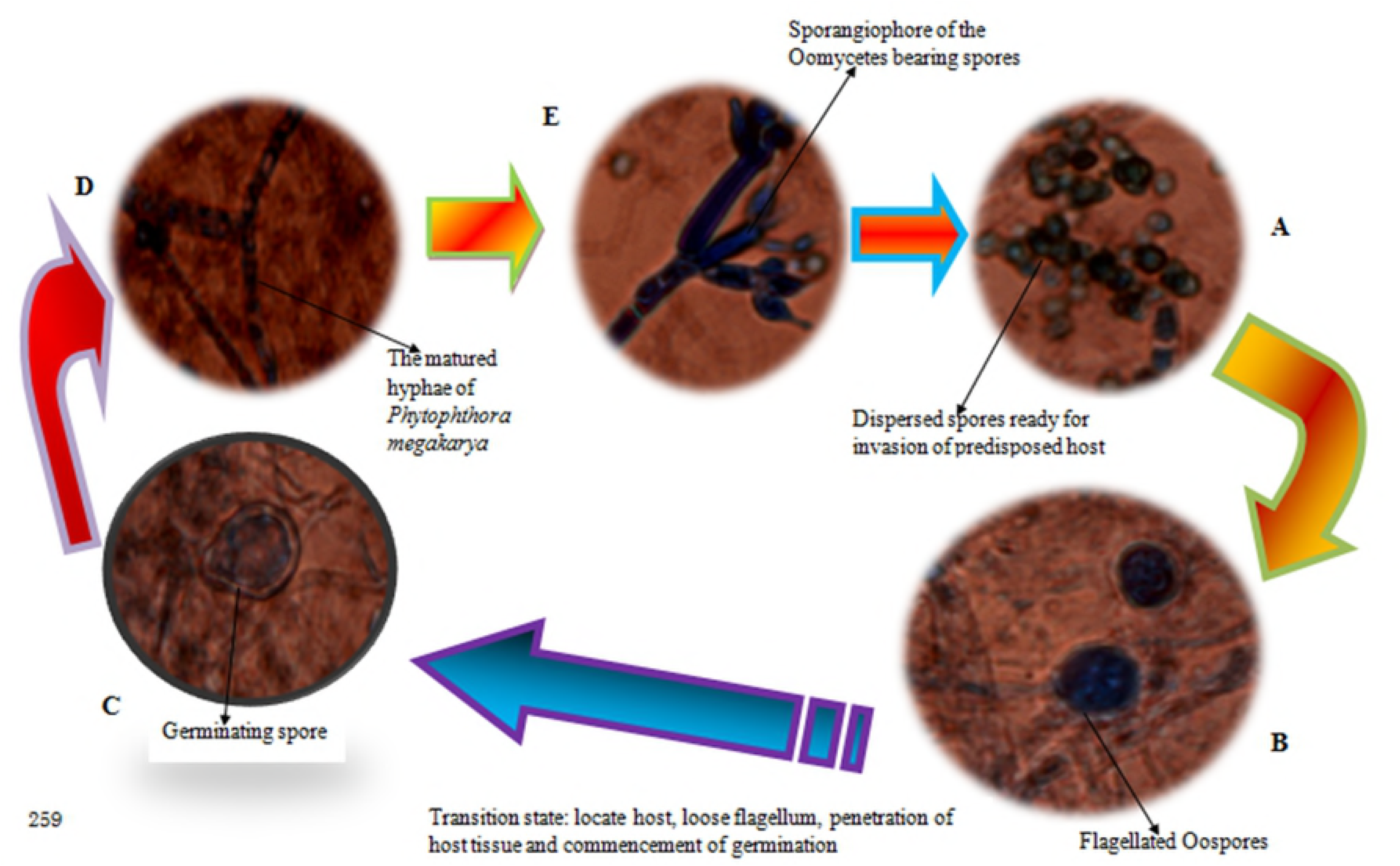
The life cycle of *Phytophthora megakarya* responsible for inciting black pod disease of cocoa in South western Nigeria.

The dispersed spores either swim towards nearby hosts based on the chemical attraction or signalling from the root rhizosphere or form thick enclosures that protect them from desiccation and other adverse environmental conditions. Spore dispersion is often aided by rain splash or insect activities (Fig 5). Pod infection start from cocoa pod closest to the soil and it is further disseminated by the activities of insects and other rodents within the cocoa field (Fig 5).

### An overview of the major Diseases/Pestilence of cocoa in South western Nigeria

The statutory disease assessment conducted in Ogun, Ondo, Osun and Oyo States in South western, Nigeria during the 2015/2016 cocoa production season showed that the probability for the period of occurrence of pests (i.e. insect activities, rodents etc.) was 0.5 (50%), with the activities of these pests severely rampant in the dry season. Cherelle wilt was noticed in all the cocoa farms assessed in the dryer periods of the year 0.5 (50%) with no traces during the wet season. It was further observed that black pod disease had the same level of probability in terms of seasonal occurrence [0.5 (50%)] but the occurrence of black pod disease was majorly prominent in the rainy season (Table 3).

**Table 3:**
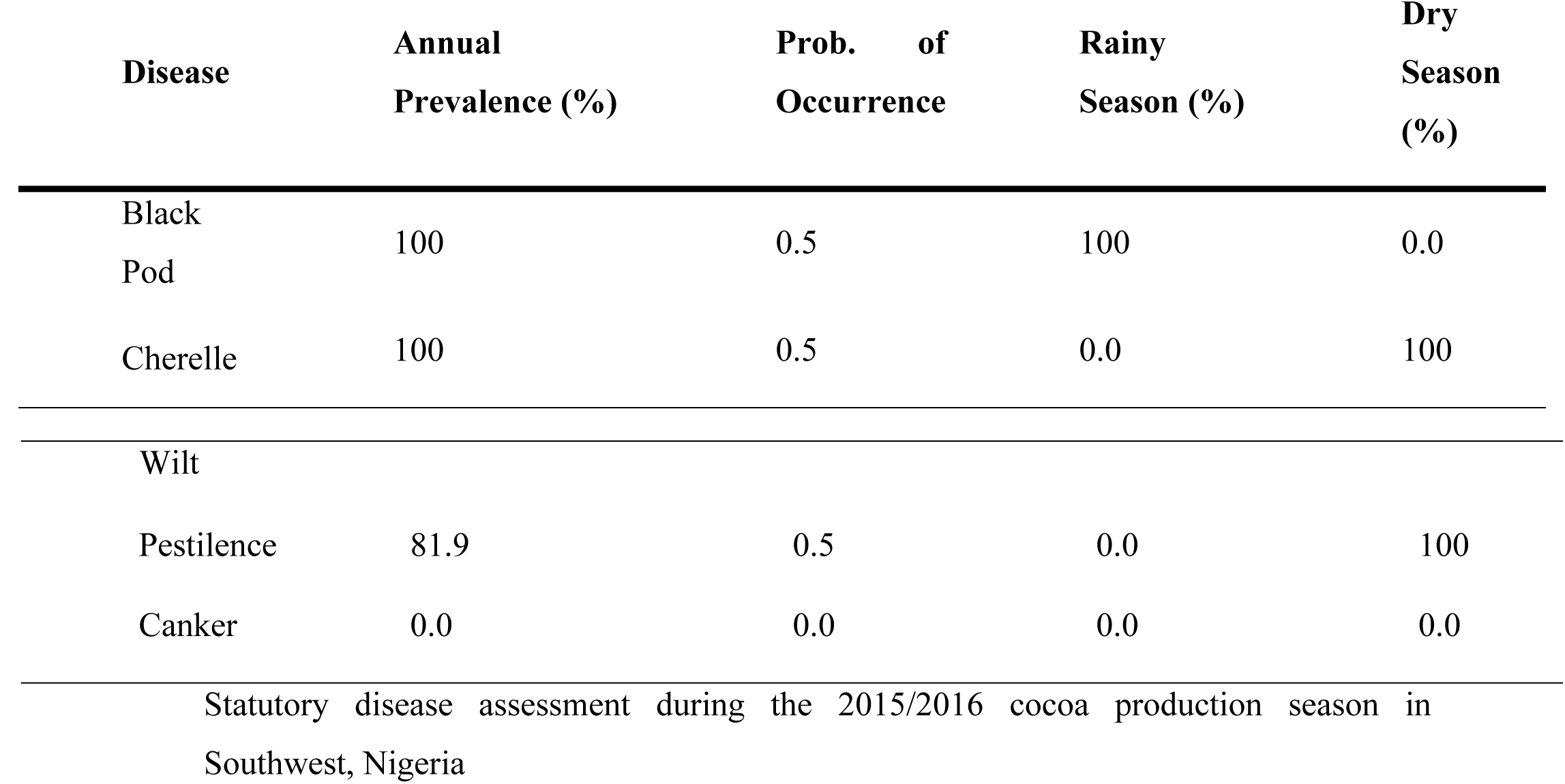
General estimation of the prevalence of diseases and pestilence of cocoa in Southwest, Nigeria

### The Level of Diseases and Pest Invasion in South western Nigeria

The level of black pod disease epidemics during the wet season was 100% across all the states. Cherelle wilt had 100% occurrence in the dry season but it was not as intensive as black pod disease. Insect and Pest invasion was 81.9% as it was undetected in some of the cocoa farms assessed (Table 3). There were no observable symptoms of stem canker in all the cocoa farms assessed from the far end of Ondo State to the rural communities in Ogun State. Therefore, the possibility of occurrence of stem canker within these regions was 0% and as such, one less problem for local cocoa farmers to contend with (Table 3).

### Black pod disease outbreak in the sampled Stations

Black pod disease (BPD) was noticed early in Stations 5 and 6 with 3.0 and 9.0% level of epidemics respectively for May 2015 (Table 4). There were no visible signs or symptoms of the disease in other Stations (0.0% BPD incidence). In June 2015, Station 3 had the highest recorded BPD occurrence (12.0%), Station 5 was second on the disease profile list with BPD outbreak of 11.0%. Stations 1, 2 and 4 had slightly severe epidemics (8.0, 7.0 and 7.0% respectively). Other Stations had NO epidemics (0.0%) as shown in Fig 6.

**Table 4:** Profile of black pod disease outbreak within the sampled stations

| Average Monthly Disease Occurrence (%) |  |  |  |  |  |  |  |  |  |  |  |  |
| --- | --- | --- | --- | --- | --- | --- | --- | --- | --- | --- | --- | --- |
| Period | Òwenà | Òwenà-2 | Wáàsimi | Adaàgbà | Iyánfowórogi | O-Igàngán | Owódé | Q-Owódé | Q-Owódé | Mòyè | D-Lógbà | Olórò |
| May 2015 | 0.0<sup>c</sup> | 0.0<sup>c</sup> | 0.0<sup>c</sup> | 0.0<sup>c</sup> | 3.0<sup>b</sup> | 9.0<sup>a</sup> | 0.0<sup>c</sup> | 0.0<sup>c</sup> | 0.0<sup>c</sup> | 0.0<sup>c</sup> | - | - |
| June 2015 | 7.0<sup>b</sup> | 8.0<sup>b</sup> | 12.0<sup>a</sup> | 7.0<sup>b</sup> | 11.0<sup>a</sup> | - | - | 0.0<sup>c</sup> | 0.0<sup>c</sup> | - | - | - |
| July 2015 | 20.0<sup>a</sup> | - | 16.0<sup>b</sup> | 12.0<sup>c</sup> | 15.0<sup>b</sup> | - | - | 0.0<sup>e</sup> | 0.0<sup>e</sup> | - | 6.0<sup>d</sup> | - |
| Aug.2015 | 30.0<sup>a</sup> | - | 23.0<sup>b</sup> | 9.0<sup>d</sup> | 7.0<sup>d</sup> | - | - | - | 3.0<sup>e</sup> | - | 16.0<sup>c</sup> | - |
| Sept.2015 | 10.0<sup>c</sup> | - | 12.0<sup>bc</sup> | 16.0<sup>a</sup> | 7.0<sup>d</sup> | - | - | - | 15.0<sup>a</sup> | - | 14.0<sup>ab</sup> | - |
| Oct. 2015 | 6.0<sup>c</sup> | 0.0<sup>e</sup> | 4.0<sup>d</sup> | 13.0<sup>b</sup> | 7.0<sup>c</sup> | 0.0<sup>e</sup> | - | - | 22.0<sup>a</sup> | 0.0<sup>e</sup> | 0.0<sup>e</sup> | 0.0<sup>e</sup> |
| Nov.2015 | 0.0<sup>a</sup> | 0.0<sup>a</sup> | 0.0<sup>a</sup> | 0.0<sup>a</sup> | 0.0<sup>a</sup> | 0.0<sup>a</sup> | - | - | 0.0<sup>a</sup> | 0.0<sup>a</sup> | 0.0<sup>a</sup> | 0.0<sup>a</sup> |
| Dec. 2015 | 0.0<sup>a</sup> | 0.0<sup>a</sup> | 0.0<sup>a</sup> | 0.0<sup>a</sup> | 0.0<sup>a</sup> | 0.0<sup>a</sup> | - | - | 0.0<sup>a</sup> | 0.0<sup>a</sup> | 0.0<sup>a</sup> | 0.0<sup>a</sup> |
| Jan. 2016 | 0.0<sup>a</sup> | 0.0<sup>a</sup> | 0.0<sup>a</sup> | 0.0<sup>a</sup> | 0.0<sup>a</sup> | 0.0<sup>a</sup> | - | - | 0.0<sup>a</sup> | 0.0<sup>a</sup> | 0.0<sup>a</sup> | 0.0<sup>a</sup> |
| Feb.2016 | 0.0<sup>a</sup> | 0.0<sup>a</sup> | 0.0<sup>a</sup> | 0.0<sup>a</sup> | 0.0<sup>a</sup> | 0.0<sup>a</sup> | - | - | 0.0<sup>a</sup> | 0.0<sup>a</sup> | 0.0<sup>a</sup> | 0.0<sup>a</sup> |
| Mar.2016 | 0.0<sup>a</sup> | 0.0<sup>a</sup> | 0.0<sup>a</sup> | 0.0<sup>a</sup> | 0.0<sup>a</sup> | 0.0<sup>a</sup> | - | - | 0.0<sup>a</sup> | 0.0<sup>a</sup> | 0.0<sup>a</sup> | 0.0<sup>a</sup> |
| Apr.2016 | 0.0<sup>a</sup> | 0.0<sup>a</sup> | 0.0<sup>a</sup> | 0.0<sup>a</sup> | 0.0<sup>a</sup> | 0.0<sup>a</sup> | - | - | 0.0<sup>a</sup> | 0.0<sup>a</sup> | 0.0<sup>a</sup> | 0.0<sup>a</sup> |
| May 2016 | 0.0<sup>a</sup> | 0.0<sup>a</sup> | 0.0<sup>a</sup> | 0.0<sup>a</sup> | 0.0<sup>a</sup> | 0.0<sup>a</sup> | - | - | 0.0<sup>a</sup> | 0.0<sup>a</sup> | 0.0<sup>a</sup> | 0.0<sup>a</sup> |

Means with the same alphabets across the row are not significantly different at P<0.05 using Duncan Multiple Range Test (DMRT) for
separation of statistically significant means. Data collected were represented as “Means” only
**O-Igàngán** : Owódé-Igàngán, **Q-Owódé** : Qbafèmi-Owódé, **D-Lógbà**: Dáagi-Lógbà

**Fig 6:**
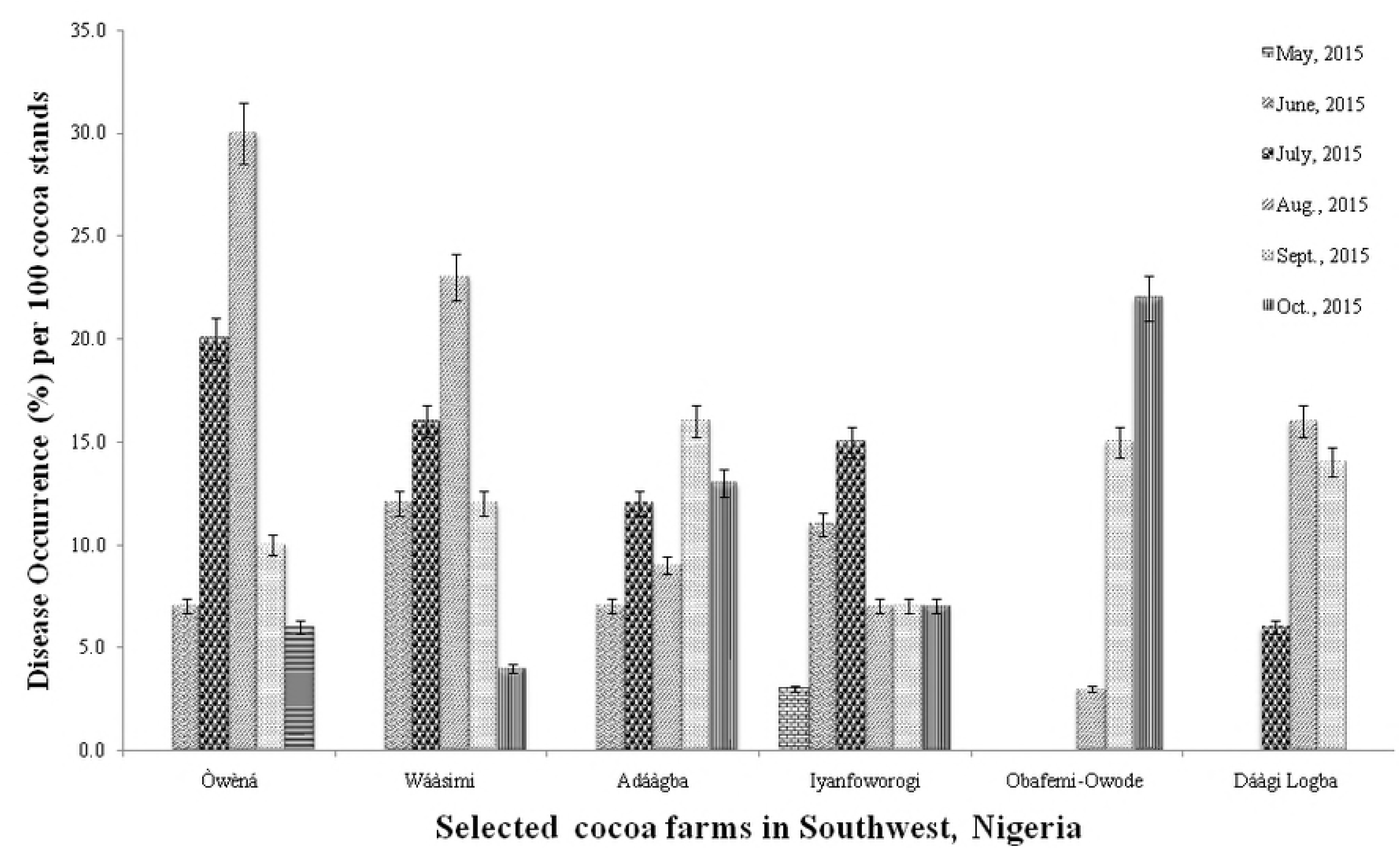
Black pod disease incidence across Southwest, Nigeria for the 2015/2016 cocoa production season

There was decline in BPD epidemics in September 2015, Station 4 took the lead (16.0%), followed by Station 8 with 15.0% BPD epidemics. Stations 11, 3, 1, and 5 had slightly severe to mild BPD outbreak of 14.0, 12.0, 10.0, and 7.0% respectively. Further decline in disease outbreak was noted in some Stations in the month of October like Station 11 (0.0%), Station 3 (4.0%), Station 1 (6.0%), Station 5 (7.0%) and Station 4 (13.0%) with Station 8 showing progressive increase in black pod diseases occurrence (22.0%) as shown in Fig 6.

There were no observable symptoms or signs associated with BPD outbreak for November and December 2015, likewise January, February, March, April and May 2016 (Table 4). This was partly due to the fact that most cocoa farmers have harvested their pods from the field coupled with the fact that there was no available moisture within the soil surface to effect *Phytophthora* spore germination and dispersion.

### Black pod disease outbreak in Ondo, Ogun, Osun and Oyo States

Black pod disease outbreak was recorded early in Osun State (1.5%) in May 2015, other States had NO BPD incidence (0.0%). Ondo and Osun States experienced slight BPD outbreak (9.5 and 9.0, respectively) in the month of June 2015. There was a progressive disease increase in Ondo State (18.0%) and Osun State (13.5%) for the month of July, 2015. Ogun and Oyo States had NO known cases of BPD outbreak for these months (Table 5).

**Table 5:** Monthly black pod disease assessment in four states within Southwest Nigeria

|  | Monthly Regional Disease Occurrence (%) per 100 trees |  |  |  |
| --- | --- | --- | --- | --- |
|  | Ondo | Osun | Ogun | Oyo |
| May 2015 | 0.0<sup>b</sup> | 1.5<sup>a</sup> | 0.0<sup>b</sup> | 0.0<sup>b</sup> |
| June 2015 | 9.5<sup>a</sup> | 9.0<sup>a</sup> | 0.0<sup>b</sup> | 0.0<sup>b</sup> |
| July 2015 | 18.0<sup>a</sup> | 13.5<sup>b</sup> | 0.0<sup>d</sup> | 6.0<sup>c</sup> |
| Aug.2015 | 26.5<sup>a</sup> | 8.0<sup>c</sup> | 3.0<sup>d</sup> | 16.0<sup>b</sup> |
| Sept.2015 | 11.0<sup>a</sup> | 11.5<sup>a</sup> | 15.0<sup>a</sup> | 14.0<sup>a</sup> |
| Oct. 2015 | 5.0<sup>c</sup> | 10.0<sup>b</sup> | 22.0<sup>a</sup> | 0.0<sup>d</sup> |
| Nov.2015 | 0.0<sup>a</sup> | 0.0<sup>a</sup> | 0.0<sup>a</sup> | 0.0<sup>a</sup> |
| Dec. 2015 | 0.0<sup>a</sup> | 0.0<sup>a</sup> | 0.0<sup>a</sup> | 0.0<sup>a</sup> |
| Jan. 2016 | 0.0<sup>a</sup> | 0.0<sup>a</sup> | 0.0<sup>a</sup> | 0.0<sup>a</sup> |
| Feb.2016 | 0.0<sup>a</sup> | 0.0<sup>a</sup> | 0.0<sup>a</sup> | 0.0<sup>a</sup> |
| Mar.2016 | 0.0<sup>a</sup> | 0.0<sup>a</sup> | 0.0<sup>a</sup> | 0.0<sup>a</sup> |
| Apr.2016 | 0.0<sup>a</sup> | 0.0<sup>a</sup> | 0.0<sup>a</sup> | 0.0<sup>a</sup> |
| May 2016 | 0.0<sup>a</sup> | 0.0<sup>a</sup> | 0.0<sup>a</sup> | 0.0<sup>a</sup> |

Means with the same alphabets across the row are not significantly different at $P < 0.05$ using Duncan Multiple Range Test (DMRT) for separation of statistically significant means. Data collected were represented as “Means” only

Ondo State maintained top position in BPD outbreak (26.5%) for August 2015, closed followed by Oyo State (16.0%). Osun State and Ogun State had 8.0% and 3.0% respectively. An increase in BPD outbreak was experienced in Ogun State from the month of September (15.0%) through October (22.0%) in 2015, whereas Osun, Ondo and Oyo State experienced decline in BPD outbreak from 11.5 to 10.0%, 11.0 to 5.0%, and 14.0 and 0.0%, respectively (Table 5). There was no incidence of black pod disease prevalence in the dry season for these States.

### Farm altitude and black pod disease Outbreak

Cocoa farms located above 200m from sea level (˃200m) experienced early BPD outbreak, beginning from May (0.8%) through August (17.3%) 2015 followed by a sharp decline from the month of September (11.3%) through October (7.5%) and 0% black pod disease incidence within the dryer period of the season. This result was an Indication of the importance of altitude in black pod disease development (Table 6).

**Table 6:**
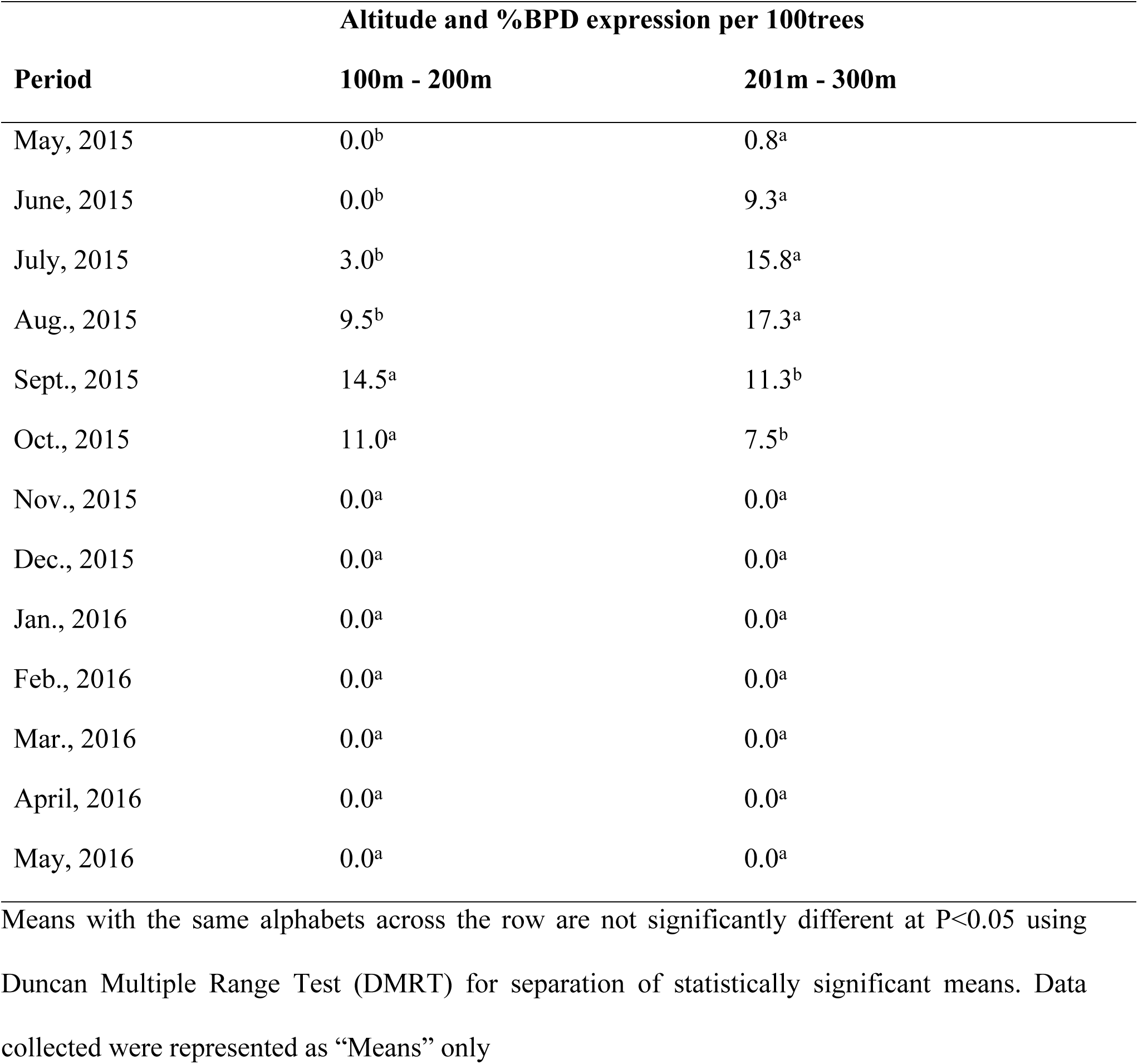
Different Altitude levels of the sampled stations and its effects on black pod disease outbreak

Cocoa farmlands located in regions that are below 200m above sea level (≤200m) had a slow start to black pod disease development (Table 6) with the peak level of black pod disease prevalence in September 2015 (14.5%) and an abrupt decline in October (11.0%) in 2015 through May 2016. A cross comparison between the two altitudes suggest that the activities of the pathogen was closely affected by the altitude of the cocoa farmlands, whereas the mode of spread was a function of the topography of the environment.

### Black pod disease prevalence and Severity in South western Nigeria

The average disease occurrence within the Southwest for the month of May 2015 was 0.4%. Further records taken for the preceding months are as follow; June 2015 (4.6%), July 2015 (9.4%), and August 2015 was the peak of disease occurrence within this terrain with an average value of 13.4% (per one hundred cocoa trees sampled). A decline in disease value was observed in the month of September 2015, with an average value of 12.9 %. Further decline in value occurred in October 2015 (9.3%) prior to harvesting of cocoa pods by farmers. The months of November, December 2015, January, February, March, April and May 2016 had negligible and unsubstantial amount of disease prevalence (Table 7). The same trend of disease report was observed within this zone for the disease severity. The peak of disease intensity was recorded in September 2015 with a mean value of 86.8%, while the least recorded occurrences were in the months of November, December 2015, January, February, March, April and May 2016 with 0.0% disease intensity (Table 7).

**Table 7:**
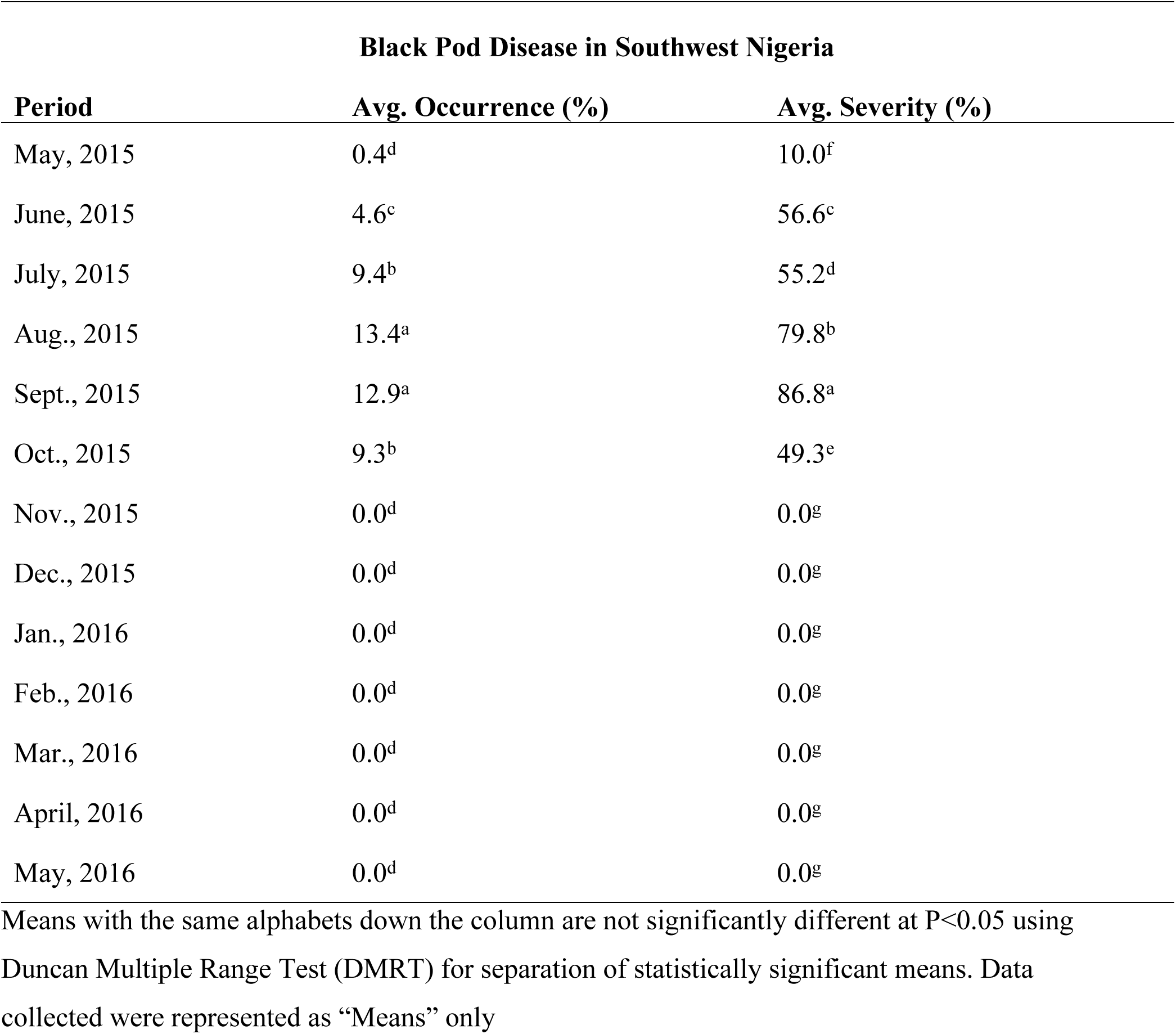
Black pod disease profile from May 2015 to May 2016 in Southwest, Nigeria

| **Black Pod Disease in Southwest Nigeria** |  |  |
| --- | --- | --- |
| **Period** | **Avg. Occurrence (%)** | **Avg. Severity (%)** |
| May, 2015 | 0.4<sup>d</sup> | 10.0<sup>f</sup> |
| June, 2015 | 4.6<sup>c</sup> | 56.6<sup>c</sup> |
| July, 2015 | 9.4<sup>b</sup> | 55.2<sup>d</sup> |
| Aug., 2015 | 13.4<sup>a</sup> | 79.8<sup>b</sup> |
| Sept., 2015 | 12.9<sup>a</sup> | 86.8<sup>a</sup> |
| Oct., 2015 | 9.3<sup>b</sup> | 49.3<sup>e</sup> |
| Nov., 2015 | 0.0<sup>d</sup> | 0.0<sup>g</sup> |
| Dec., 2015 | 0.0<sup>d</sup> | 0.0<sup>g</sup> |
| Jan., 2016 | 0.0<sup>d</sup> | 0.0<sup>g</sup> |
| Feb., 2016 | 0.0<sup>d</sup> | 0.0<sup>g</sup> |
| Mar., 2016 | 0.0<sup>d</sup> | 0.0<sup>g</sup> |
| April, 2016 | 0.0<sup>d</sup> | 0.0<sup>g</sup> |
| May, 2016 | 0.0<sup>d</sup> | 0.0<sup>g</sup> |

Means with the same alphabets down the column are not significantly different at P<0.05 using Duncan Multiple Range Test (DMRT) for separation of statistically significant means. Data collected were represented as “Means” only

### Black pod disease profile in Ogun, Ondo, Osun and Oyo States (Southwest, Nigeria)

The general field assessment conducted in 2015/2016 showed that Ondo State had the highest level of black pod disease prevalence (Incidence) and Osun State had the highest level of disease Intensity (Severity) with average annual values of 8.8% for BPD outbreak (Ondo) and 53.1% (Osun) for BPD severity (Table 8). The general result for BPD outbreak and prevalence was recorded thus: Ondo State (8.8%, 46.2%), Osun State (6.7%, 53.1%), Ogun State (5.0%, 31.7%), and Oyo State (4.5%, 39.3%) as reported in Table 8.

**Table 8:** Seasonal Changes and its

| State | Occurrence (%) |  | Severity (%) |  |
| --- | --- | --- | --- | --- |
|  | Rainy Season | Dry Season | Rainy Season | Dry Season |
| Ondo | 8.8<sup>a</sup> | 0.0<sup>a</sup> | 46.2<sup>ab</sup> | 0.0<sup>a</sup> |
| Osun | 6.7<sup>b</sup> | 0.0<sup>a</sup> | 53.1<sup>a</sup> | 0.0<sup>a</sup> |
| Ogun | 5.0<sup>bc</sup> | 0.0<sup>a</sup> | 31.7<sup>c</sup> | 0.0<sup>a</sup> |
| Oyo | 4.5<sup>c</sup> | 0.0<sup>a</sup> | 39.3<sup>bc</sup> | 0.0<sup>a</sup> |

Means with the same alphabets down the column are not significantly different at $P < 0.05$ using Duncan Multiple Range Test (DMRT) for separation of statistically significant means. Data collected were represented as “Means” only.

The overall annual estimation of BPD intensity and outbreak within Southwest, Nigeria (2015/2016) was 42.6% and 6.2% respectively (Table 9).

**Table 9:** The average black pod disease status in Southwest, Nigeria (2015/2016)

| **Black Pod Disease** | **Rainy Season (%)** | **Dry Season (%)** |
| --- | --- | --- |
| **Occurrence** | 6.2<sup>a</sup> | 0.0<sup>b</sup> |
| **Severity** | 42.6<sup>a</sup> | 0.0<sup>b</sup> |

Means with the same alphabets across the row are not significantly different at $P < 0.05$ using Duncan Multiple Range Test (DMRT) for separation of statistically significant means. Data collected were represented as “Means” only

### Black pod disease severity in the sampled Stations

Black pod disease was expressed early in Stations 5 and 6 (60.0% and 71.1%). Other Stations had 0.0% disease expression in the month of May, 2015. The disease severity for June 2015 was 93.9% for Station 3, 62.5% for Station 2, 60.0% for Station 1, 95.1% for Station 4, and 90.9% for Station 5. Other Stations had 0.0% BPD severity within that period (Table 10). Station 1 had the highest black pod disease intensity in the Southwest (87.5%) in the month of July, closely followed by Station 4 (78.1%), Station 11 (76.7%), Station 5 (62.2%), and Station 3 (60.0%) respectively.

**Table 10:**
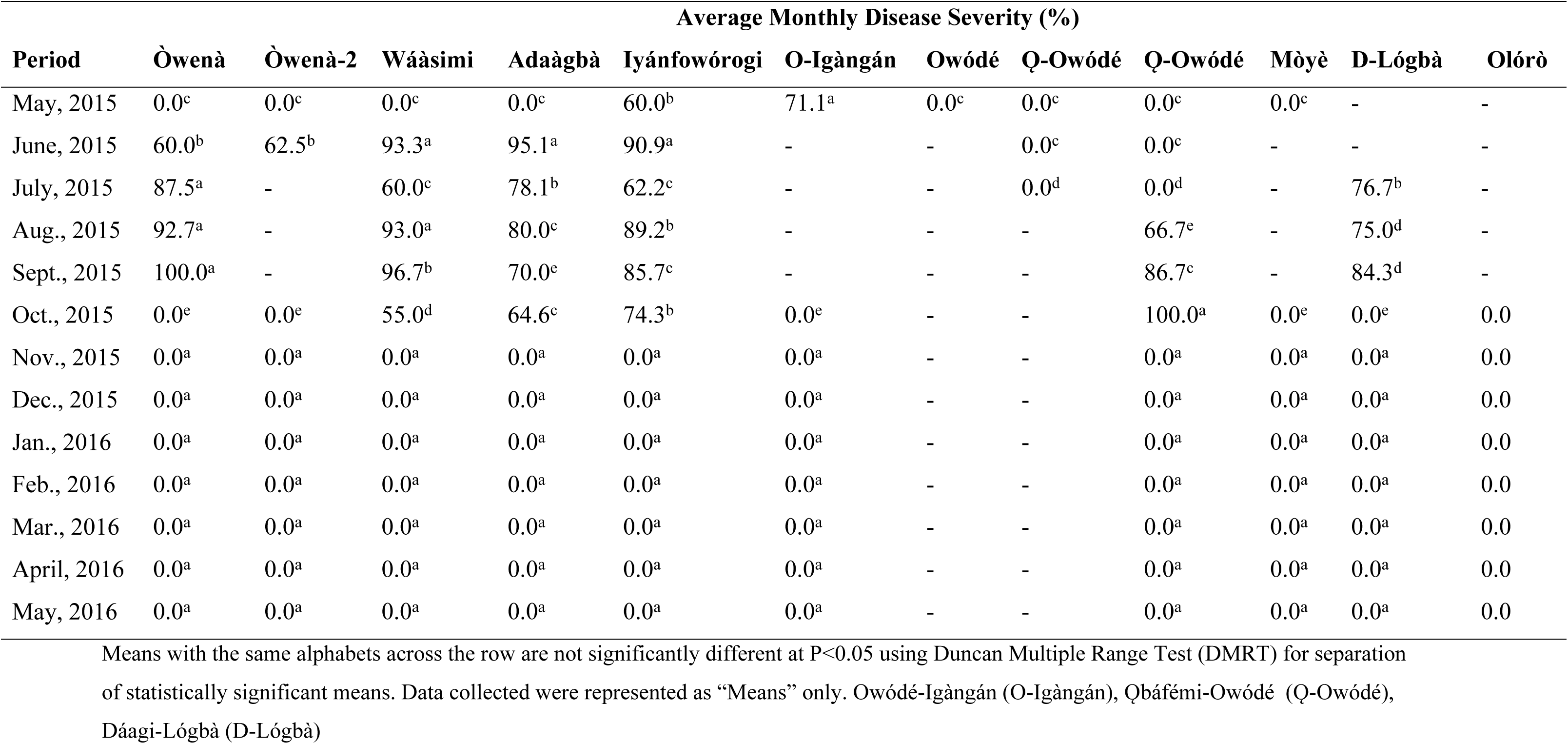
Black pod disease severity levels within the selected stations

In August, Station 3 had the highest BPD severity level (93.0%) in the Southwest of Nigeria, closely followed by Station 1 (92.7%), Station 5 (89.2%), Station 4 (80.0%), Station 11 (75.0%), and Station 8 (66.7%) as documented in Table 12. In September 2015, Station 1 had 100.0% BPD severity, Station 3 had 96.7%, Station 8 had 86.7%, Station 5 had 85.7%, Station 11 had 84.3%, and Station was the least with 70.0%. There was a massive decline in disease outbreak in some locations in the month of October like Station 11 (0.0%), Station 3 (55.0%), Station 1 (0.0%). Stations 4, 5, and 8 still had high BPD severity with the following values 64.6, 74.3 and 100.0%, respectively (Table 10). There were no observable symptoms of BPD within in the later periods of the season.

### Black pod disease intensity in Ondo, Ogun, Osun and Oyo States

The level of black pod disease severity was milder in the early periods of the 2015/2016 cocoa production season from the statutory disease assessment result documented in Table 11. It was observed that the intensity of the disease was more in Osun state with mean disease severity of 30.0% in the month of May 2015 (during the rainy season), whereas, other states within the Southwest had no signs or symptoms of the disease (0.0% BPD severity). The persistency of black pod disease was more in June across cocoa farms located in Osun State (93.0%) and Ondo State (76.7%). The intensity of black pod disease was relatively insignificant in other States sampled within the same period.

**Table 11:** An evaluation of the average level of black pod disease severity in some Southwestern States

| Period | Black Pod Disease Severity (%) |  |  |  |
| --- | --- | --- | --- | --- |
|  | Ondo | Osun | Ogun | Oyo |
| May, 2015 | 0.0<sup>b</sup> | 30.0<sup>a</sup> | 0.0<sup>b</sup> | - |
| June, 2015 | 76.7<sup>b</sup> | 93.0<sup>a</sup> | 0.0<sup>c</sup> | - |
| July, 2015 | 73.8<sup>b</sup> | 70.1<sup>c</sup> | 0.0<sup>d</sup> | 76.7<sup>a</sup> |
| Aug., 2015 | 92.9<sup>a</sup> | 84.6<sup>b</sup> | 66.7<sup>d</sup> | 75.0<sup>c</sup> |
| Sept., 2015 | 98.3<sup>a</sup> | 77.9<sup>d</sup> | 86.7<sup>b</sup> | 84.3<sup>c</sup> |
| Oct., 2015 | 27.5<sup>c</sup> | 69.5<sup>b</sup> | 100.0<sup>a</sup> | 0.0<sup>d</sup> |
| Nov., 2015 | 0.0<sup>a</sup> | 0.0<sup>a</sup> | 0.0<sup>a</sup> | 0.0<sup>a</sup> |
| Dec., 2015 | 0.0<sup>a</sup> | 0.0<sup>a</sup> | 0.0<sup>a</sup> | 0.0<sup>a</sup> |
| Jan., 2016 | 0.0<sup>a</sup> | 0.0<sup>a</sup> | 0.0<sup>a</sup> | 0.0<sup>a</sup> |
| Feb., 2016 | 0.0<sup>a</sup> | 0.0<sup>a</sup> | 0.0<sup>a</sup> | 0.0<sup>a</sup> |
| Mar., 2016 | 0.0<sup>a</sup> | 0.0<sup>a</sup> | 0.0<sup>a</sup> | 0.0<sup>a</sup> |
| April, 2016 | 0.0<sup>a</sup> | 0.0<sup>a</sup> | 0.0<sup>a</sup> | 0.0<sup>a</sup> |
| May, 2016 | 0.0<sup>a</sup> | 0.0<sup>a</sup> | 0.0<sup>a</sup> | 0.0<sup>a</sup> |

Means with the same alphabets across the row are not significantly different at $P < 0.05$ using Duncan Multiple Range Test (DMRT) for separation of statistically significant means. Data collected were represented as “Means” only

There was rapid geometric increase in black pod disease expression and intensity within the months of July and August with a climax in September 2015 (Table 11). The disease severity for July was recorded thus: Ondo State (73.8%), Osun State (70.1%), Oyo State (76.7%) and Ogun State (0.0%). In August, Ondo State recorded 92.9% BPD intensity; Osun State had 84.6%, Ogun State (66.7%) and Oyo State (75.0%).

The disease intensity across Ogun, Ondo, Osun and Oyo States during the 2015/2016 cocoa production season for September 2015 was 98.3% (Ondo State), 86.7% (Ogun State), 84.3% (Oyo State) and 77.9% (Osun State). Ogun State had 100.0% disease intensity in the month of October 2015 which was contrary to the trend of disease progress in other States. Ondo State had 27.5%, Osun State (69.5%) and Oyo State (0.0%). Subsequently, other months within the dryer periods of the year (2015/2016 cocoa production season) had insignificant disease intensity values (0%). The findings were recorded in Table 11.

### Altitude and its effects on black pod disease expression

It was observed that 15.0% black pod disease intensity was recorded for the month of May 2015 for cocoa farmlands located at altitude higher than 200m above sea level (201m-300m) and 0.0% for cocoa plantations located within or below 200m above sea level (Table 12). The same trend was observed in June 2015 with 84.8% black pod disease intensity for cocoa farmlands situated within altitude 201m-300m and 0.0% for those located within or below 200m.

**Table 12:** The influence of altitude on black pod disease severity in Southwest Nigeria

| **Duration** | **Height Above Sea Level** |  |
| --- | --- | --- |
|  | **100m - 200m** | **201m - 300m** |
| May, 2015 | 0.0<sup>b</sup> | 15.0<sup>a</sup> |
| June, 2015 | 0.0<sup>b</sup> | 84.8<sup>a</sup> |
| July, 2015 | 38.4<sup>b</sup> | 71.9<sup>a</sup> |
| Aug., 2015 | 70.9<sup>b</sup> | 88.7<sup>a</sup> |
| Sept., 2015 | 85.5<sup>a</sup> | 88.1<sup>b</sup> |
| Oct., 2015 | 50.0<sup>a</sup> | 48.5<sup>a</sup> |
| Nov., 2015 | 0.0<sup>a</sup> | 0.0<sup>a</sup> |
| Dec., 2015 | 0.0<sup>a</sup> | 0.0<sup>a</sup> |
| Jan., 2016 | 0.0<sup>a</sup> | 0.0<sup>a</sup> |
| Feb., 2016 | 0.0<sup>a</sup> | 0.0<sup>a</sup> |
| Mar., 2016 | 0.0<sup>a</sup> | 0.0<sup>a</sup> |
| April, 2016 | 0.0<sup>a</sup> | 0.0<sup>a</sup> |
| May, 2016 | 0.0<sup>a</sup> | 0.0<sup>a</sup> |

BPD intensity was progressive through the months of July 2015 (38.4% for cocoa farmlands located below 200m altitude, and 71.9% for cocoa plantations above 200m), August 2015 (70.9% for cocoa fields below 200m altitude, and 88.7% for areas above 200m), with its peak value in September 2015 for cocoa farmlands located below 200m height above sea level (85.5% black pod disease expression), and 88.1% for areas above 200m in altitude. There was a decline in black pod disease expression in October 2015, with a recorded value of 50.0% for areas situated below or within 200m height above sea level and 48.5% for areas above 200m. There was no expression of the disease during the dry season (Table 12).

## Discussion

### Black pod disease diagnosis in Ogun, Ondo, Osun and Oyo States, Nigeria

It was evident from the statutory disease assessment conducted that black pod disease was a major threat to cocoa production in Southwestern Nigeria. Its occurrence was recurrent and rapid with massive destruction and devastating effects as it spreads. Some areas identified to be under severe black pod disease attack include Òwenà and Wáàsimi in Ondo East local government area (LGA) of Ondo State, Adaàgbà and Iyánfowórogi in Ife South LGA of Osun State, Owódé- Igàngán and its’ environs in Àtàkúnmòsà East LGA of Osun State, Ǫbáfémi-Owódé in Abeokuta (Ogun State), Dáagi-Lógbà and Atérè Villages within Ǫmi-Adió area of Iddo LGA, Olórò Village (Olorunda) in Àkànràn, and Mòyè village in Ǫnà-Arà LGA of Oyo State. This report was a confirmation of the findings of Adisa and Adeloye [25] who stated that these areas (particularly Osun State) have been identified as hotspots for black pod disease invasion, urgently in need of integrated disease management strategies.

### The trend of black pod disease occurrence in Southwest, Nigeria

It was noted that the severity of black pod disease, its prevalence and spread was widely pronounced in the rainy season and there was 100% chances of black pod disease occurrence in all the cocoa farmlands investigated as long as there was consistent pattern of rainfall within that terrain. This was in agreement with the observations made by Adeniyi and Ogunsola [26] that the yearly variation in the yield of cocoa is affected more by rainfall and the disease thrives better when there is moisture available in the surrounding environment. Finally, the observations made by Anim-Kwapong and Frimpong [27] is a suggestion that cocoa is highly sensitive to changes in weather pattern which can greatly influence its yield and productivity, and predispose it to a greater extent to black pod disease infection.

Other findings accrued from the research conducted showed that black pod disease was the most influential among all the diseases limiting cocoa production in Southwest, Nigeria. It was also noticed that insects and pests attack on green and ripe cocoa pods occurred during the dry season at epidemic proportions. Akrofi [24] consented to this fact but he did not give any information on the season of occurrence of these pests. A combination of the multiple effects of the pests and diseases was a major threat to local cocoa farmers due to their high tendency to cause yield reduction, their fast spreading nature in the field and high devastating effects in terms of pod destruction, depletion in quality nutrients of the bean, distortion of cocoa bean texture, colour and reduction of market values made it a major challenge for local cocoa growers within these terrain. A similar observation was also recorded by Flood co-workers [28].

### Black pod disease profile in the rural areas of Southwest, Nigeria

Black pod disease was noticed early in “Owódé-Igàngán”, an area close to Ilèsà within Àtàkúnmòsà East of Osun State and “Iyánfowórogi” located in Ife South (both in Osun State) in the month of May, 2015; few months after the commencement of the rainy season. This was indeed a reaffirmation of the statement made by Adisa and Adeloye [25] that Osun State has been identified as a black pod disease prone region. Other research locations had no prevalence of the disease. This was partly due to the fact that the environment was not favourable enough at that point in time for the proliferation of black pod disease pathogen.

There was a change in the disease trend for the month of June 2015 as “Wáàsimi” (an area located in the heart of Ondo State) had the highest black pod disease occurrence, closely followed by “Iyánfowórogi” in Ife South, “Òwenà (still in Ondo), and Adaàgbà (a village in close proximity with “Iyánfowórogi” in Ife South of Osun State). Other locations like Ogun State and Oyo State were black pod disease free. The same pattern of disease trend was described by Opoku [3] in Ghana that the primary infections usually occur around June, but the peak of black pod disease invasion generally occurs between August and October.

A similar pattern of disease spread was also observed for the month of July but in this case “Òwenà and Wáàsimi had massive black pod disease incidence. Iyánfowórogi and Adaàgbà too had substantial disease spread which was becoming a cause for worry to the local farmers. Dáagi-Lógbà in Ǫnà-Arà local Government of Oyo State surprisingly joined the group of black pod disease infected areas. This was still in concordance with previous findings [3, 7].

The pinnacle of infection was in August 2015 where the disease prevalence was at its peak, unfortunately the same trend of black pod disease spread persisted, Òwenà and Wáàsimi (both leading cocoa growing communities not just in the Southwest but in Nigeria at large) were at the forefront of the disease predicament and as such the spotlight for research focus. Infected areas like these with massive input in commercial cocoa production are regarded by epidemiologists as the “area under disease curve” (AUDC). Farmlands in areas like Dáagi-Lógbà, Adaàgbà, Iyánfowórogi and Ǫbáfémi-Owódé also had their fair share of infection. A similar observation was also noted by Appiah [7] in the tropical regions of Ghana (where cocoa are widely cultivated) that the height of *P. megakarya* induced black pod disease infection generally occurs between August and October.

In September 2015, there was a change in disease preference with Adaàgbà taking the lead trailed by Ǫbáfémi-Owódé, Dáagi-Lógbà, Wáàsimi, Òwenà, and Iyánfowórogi with the least level of black pod disease prevalence. There was a massive decline in disease outbreak in some locations in the month of October like Dáagi-Lógbà, Wáàsimi and Òwenà, Iyánfowórogi and Adaàgbà etc. Interestingly, Ǫbáfémi-Owódé showed a progressive geometric increase in black pod diseases occurrence. The observations made from other locations (with the exception of Ǫbáfémi-Owódé) were in contrast with the assertions given by Opoku [29] who noted that the peak period for black pod disease invasion were between the months of August and October where there is sufficient amount of surface water in the surrounding environment and high pathological activities of the pathogen.

With the onset of the dry season, culminating in a drastic reduction in the top soil surface water, reduced amount of rainfall, high ambient temperature, increased hours of sunshine (high luminous potentials), decreased air saturation (low relative humidity) and due to the fact that most cocoa farmers have harvested their pods from the farm, black pod disease occurrence for the months of November and December 2015, likewise January, February, March, April and May 2016 was drastically reduced to non-significant levels, and as such the disease posed no threats to farmers during these period. This was in agreement with the reports given by Oluyole and Lawal [30] who reiterated that black pod disease is prevalent only during the wet season.

### Black pod disease profile in Ogun, Ondo, Osun and Oyo States

The statutory investigation of the average monthly black pod disease prevalence for Ondo, Ogun, Osun and Oyo States was such that Osun State had a significant level of occurrence for the month of May 2015 compared to other States with no recorded history or field reports for black pod disease prevalence in May. Ondo and Osun States had a seemingly similar level of disease prevalence, co-occupying the top position for black pod disease spread during the month of June 2015, whereas there was no report of the disease in Ogun and Oyo States. This can be partly explained by the similarity in weather pattern based on proximity in distance and agro-ecological zoning of these States, a theme well described by Ziervogel [31] who stated that climate change has wide-ranging effects on the environment including water resources, agriculture, food security, human health, terrestrial ecosystems, biodiversity and coastal zones.

In July 2015, Ondo State had the highest level of black pod disease occurrence, seconded by Osun State. Other States had little, insignificant or no disease prevalence at the point of investigation. The preference for disease spread was uttered in the month of August 2015 with Ondo State maintaining top position, subsequently followed by Oyo State. Osun State and Ogun State just had minimal levels of black pod disease prevalence. Surprisingly, Ogun State had the highest values for disease prevalence for the months of September and October 2015 during the statutory disease assessment routine, followed by Oyo State, Osun State and lastly, Ondo State. There was no occurrence of black pod disease in the dry season for all the cocoa farms investigated in the course of this research. This was totally due to the fact that there was no surface water to facilitate the proliferation and spread of the pathogen and most cocoa farmers had harvested all their cocoa pods from the field. This was in agreement with the reports given by Oluyole and Lawal [30] who reiterated that black pod disease is prevalent only during the rainy season and that moisture is a pertinent factor for the development and spread of the disease.

### Topography of cocoa farmland and its influence on black pod disease development

It was learned in connection with this research that the establishment, spread and prevalence of black pod disease on farmlands located in areas of high altitude (˃200m) had geometric increase from the months of May through August 2015, followed by a rapid decline in the month of September and October. The aim of this assessment was to further determine the significance of altitude in black pod disease development. Farmlands located in regions with lesser altitude (<200m) had a slow start to black pod disease development with the pinnacle of black pod disease spread in September 2015 and an abrupt decline in October 2015 through May 2016. A cross comparison between the two height levels showed that the activities of the pathogen was closely affected by the height above sea level of the environment and the rate of spread of the disease was largely influenced to a great extent by the topography of the farmland.

### The intensity of black pod disease in Southwest Nigeria

It was observed that the trend of black pod disease development and spread in Southwest, Nigeria was logarithmical with a very slow start in the month of May 2015 and a gradual increase through the months of June 2015, through July 2015 and August 2015 were the peak of disease occurrence within the earmarked research locations were observed. A decline in black pod disease value was observed in the month of September 2015, and further decline in disease value occurred in October 2015 prior to harvesting of cocoa pods by farmers. The months of November, December 2015, January, February, March, April and May 2016 had unsubstantial amount of black disease severity.

The same trend of disease history was observed within this zone for the disease severity. The zenith of disease intensity was in September 2015 and the least recorded degrees of black pod disease intensity were for the months of November, December 2015, January, February, March, April and May 2016 with insignificant disease intensity, which was largely due to the fact that most farmers have harvested their cocoa pods and the Cherelles (young pods) which are still in the juvenile stage were not the preferential target of *Phytophthora megakarya*. During these periods the farm environment was devoid of water which is a pertinent factor for the proliferation of the organism (*Phytophthoramegakarya*) that caused black pod disease.

### The role of seasonal variation in black pod disease expression in Southwest, Nigeria

It was reported that Ondo State had the highest level of black pod disease prevalence (Incidence) during the close of the year 2015 and Osun State had the highest level of disease Intensity (Severity). These values were recorded during the rainy season. This was in agreement with the research work of Akrofi [24] who reiterated that the survival and proliferation of *Phytophthora megakarya* depends majorly on the availability of water most especially during the rainy season where water is present in abundance in most cocoa farmlands around the African continent.

### A general assessment of the development and severity of black pod disease in Southwest, Nigeria

The overall annual disease occurrence observed within Southwest, Nigeria for the cocoa production season ending 2016 were mildly severe while the levels of disease intensity on the affected cocoa pods were moderately and extremely severe in some cases. The irregular black pod disease management rate achieved around the investigated region was due to the fact that the level of preparedness of the farmers within the affected region in terms of fungicide application and good cultural practices to wade of potential agents of propagation and spread of the disease differs greatly, partly due to ignorance and the level of information on the control of the disease available to the local cocoa farmers and majorly due to financial constraints. This was in line with the findings of the Cocoa Research Institute of Nigeria (2003).

### Level of destruction of black pod disease in rural and sub-urban community of Southwest, Nigeria

The routine disease assessment conducted within twelve (12) selected rural and sub-urban settlements in Nigeria reputed for cocoa farming in the Southwest showed that the level of black pod disease intensity during the month of May 2015 was relatively high for “Owódé-Igàngán” and “Iyánfowórogi” which were regarded as key cocoa production areas in Osun State. Other locations across the Southwest of Nigeria had insignificant levels of black pod disease severity. For the month of June 2015, “Wáàsimi”, “Iyánfowórogi”, “Òwenà etc. all had significant levels of black pod disease severity, fascinatingly, Adaàgbà had the highest record of disease intensity during this period. Other locations showed no signs of black pod disease severity.

The same pattern of disease severity was also observed for the month of July but in this case Òwenà had the highest black pod disease incidence, with other sampled points like Wáàsimi, Iyánfowórogi, Adaàgbà and Dáagi-Lógbà trailing closely behind. The pinnacle of black pod disease severity was in September and not in August 2015 as conveyed in the earlier report of its spread and prevalence. It was observed that a similar trend of disease severity occurred in August within this zone, with Òwenà having a massive black pod disease infestation, Wáàsimi, Dáagi-Lógbà, Adaàgbà, Iyánfowórogi and Ǫbáfémi-Owódé too had their fair share of black pod disease infestation. This was described in pevious reports [3,7].

In September 2015, there was a change in disease preference with Òwenà having the highest mean recorded value for disease severity, Adaàgbà and other regions were reported to have severe black pod disease intensity (Ǫbáfémi-Owódé, Dáagi-Lógbà, Wáàsimi, and Iyánfowórogi). There was a massive decline in disease severity in some locations in the month of October like Dáagi-Lógbà, Wáàsimi, Òwenà, Iyánfowórogi and Adaàgbà, but surprisingly, Ǫbáfémi-Owódé showed a progressive geometric increase in black pod diseases intensity. The basic rationale behind the heavy infestation by black pod disease was in the lack of disease management strategy employed by the farmer. This confirms the assertion stated by Berry and Cilas [32] that losses can reach up to 100% of the cocoa production in smallholders’ plantations when no control measures are taken.

With the onset of the dry season, culminating in a drastic reduction in the top soil surface water, reduced amount of rainfall, high ambient temperature, increased hours of sunshine (high luminous potentials), decreased air saturation (low relative humidity) and due to the fact that most cocoa farmers have harvested their pods from the farm, black pod disease severity for the months of November and December 2015, likewise January, February, March, April and May 2016 was drastically reduced to non-significant levels, and as such the disease posed no threats to farmers during these periods. This was in line with the assertion made by Ziervogel *et al*. [31].

### Black pod disease intensity in Ogun, Ondo, Osun and Oyo States

It was observed that black pod disease started very early in cocoa farmlands located in Osun state in the month of May 2015 during the rainy season. Osun State still led the chart for high ranking black pod disease severity in the southwest for the month of June 2015, followed by Ondo State. Other States had insignificant disease history for the month of June 2015. There was rapid geometric increase in black pod disease expression and intensity within the months of July and August with a climax in September 2015 for Ondo State, Osun State, Oyo State and Ogun State. The same sequence for disease prevalence was earlier given by Appiah [7]. Other preceding months had insignificant disease intensity status.

### Landscape and black pod disease expression on cocoa pods in Nigeria

There was disease infestation recorded for cocoa farmlands located within areas situated in altitude higher than 200m above sea level (201m-300m) and none for areas located below 200m above sea level in the month of May 2015. The same trend was observed in June 2015. The disease intensity trend was progressive through the months of July 2015 and August, with its peak value in September 2015 for cocoa farmlands located in areas situated below 200m height above sea level, follow by retrogression in disease intensity value in October 2015, through the dry season. This was as suggested by Appiah [7].

## Supporting information

S1 Appendix: Details of study sites

S1 Fig: A semi-biased mode of disease sampling within an earmarked cocoa field (T - Cocoa tree)

## Conclusion

It was established by the research conducted that black pod disease was and still is a major threat to cocoa farmers in Southwest, Nigeria. Farmers within the rural communities are indeed tired of the huge loss incurred from the spread of the disease due to the recurrent and devastating effects of the disease, cocoa farming is fast becoming a myth in this region. Unless concerted efforts towards black pod disease management is put in place, cocoa farming will go into extinction in Nigeria.

